# SapBase (Sapinaceae Genomic DataBase): a central portal for functional and comparative genomics of Sapindaceae species

**DOI:** 10.1101/2022.11.25.517904

**Authors:** Jiawei Li, Chengjie Chen, Zaohai Zeng, Fengqi Wu, Junting Feng, Bo Liu, Yingxiao Mai, Xinyi Chu, Wanchun Wei, Xin Li, Yanyang Liang, YuanLong Liu, Jing Xu, Rui Xia

**Affiliations:** State Key Laboratory for Conservation and Utilization of Subtropical Agro-Bioresources, College of Horticulture, South China Agricultural University, Guangzhou, Guangdong, 510640, China; Guangdong Laboratory for Lingnan Modern Agriculture, South China Agricultural University, Guangzhou, Guangdong, 510640, China; Key Laboratory of Biology and Germplasm Enhancement of Horticultural Crops in South China, Ministry of Agriculture and Rural Affairs, South China Agricultural University, Guangzhou, Guangdong 510640, China

## Abstract

Sapindaceae is a family of flowering plants, also known as the soapberry family, comprising 141 genera and about 1900 species (Pedro et al., 2010). Most of them are distributed in tropical and subtropical regions, including trees, shrubs, also woody or herbaceous vines. Some are dioecious, while others are monoecious. Many Sapindaceae species possess great economic value; some furnish delicious fruits, like lychee (*Litchi chinensis*), longan (*Dimocarpus longan*), rambutan (*Nephelium lappaceum*); and ackee (*Blighia sapida*) - the national fruit of Jamaica; some produce abundance secondary metabolites, like saponin from soapberry (*Sapindus mukorossi*), and seed oil from yellowhorn (*Xanthoceras sorbifolium*); some yield valuable timber including maple (*Acer spp*.) and buckeye (*Aesculus glabra*); and some are of great herbal medicinal value, like balloon-vine (*Cardiospermum halicacabum*).

In the last decade, with the rocketing of next generation sequencing (NGS) and genomic technologies, the full genome sequences of several Sapindaceae plants have been resolved (Lin et al., 2017; Liang et al., 2019; Yang et al., 2019; Zhang et al., 2021; Hu et al., 2022; Xue et al., 2022). Among them, our recent publication of the lychee genome attracted broad attention (Edger, 2022; Hu et al., 2022; Lyu, 2022). Now the post-genome era arrives for Sapindaceae, however, there is no public genomic database available for any Sapindaceae species, let alone an integrative database for the whole Sapindaceae family. A unified data platform is in urgent need to collect, manage and share relevant data resources. Therefore, we integrated our home-brew NGS data with all publicly available data for seven Sapindaceae plants and constructed the **<u>Sap</u>**inaceae Genomic Data**<u>Base</u>**, named SapBase (www.sapindaceae.com), in order to provide genomic resources and an online powerful analytic platform for scientific research on Sapinaceae species and comparative studies with other plants.

## Data Source

Currently, SapBase hosts genomic resources for seven Sapindaceae species (Fig. 1A), including 16 full genome sequences, 411 sets of resequencing genomic data (411 sets, 4.82 TB), 919 RNAseq data (10.3 TB) from 49 projects, and 501 sRNA loci from the sRNAanno database (Chen et al., 2021). In total, there are 514,422 genes (893,747 transcripts), with 501,479 of them having functional annotations. 4,577 functional domains are annotated from 392,123 genes. SapBase also predicts 79,862,416 interaction relations between 145,248 proteins. 89,025 synteny blocks between every two Sapindaece species were identified covering 134,016 genes. Besides, 486 gene co-expression modules were singled out by the integrative analyses of these omics data. All these data are accessible at SapBase via the four major function categories (Fig. 1B): **BROWSE** for data and result browsing; **SEARCH** for comprehensive and efficient information retrieval; **ANALYSES** providing various data processing, analysis and visualization functions, and **DOWNLOAD** responsible for data deposit and download.

**Figure 1.**
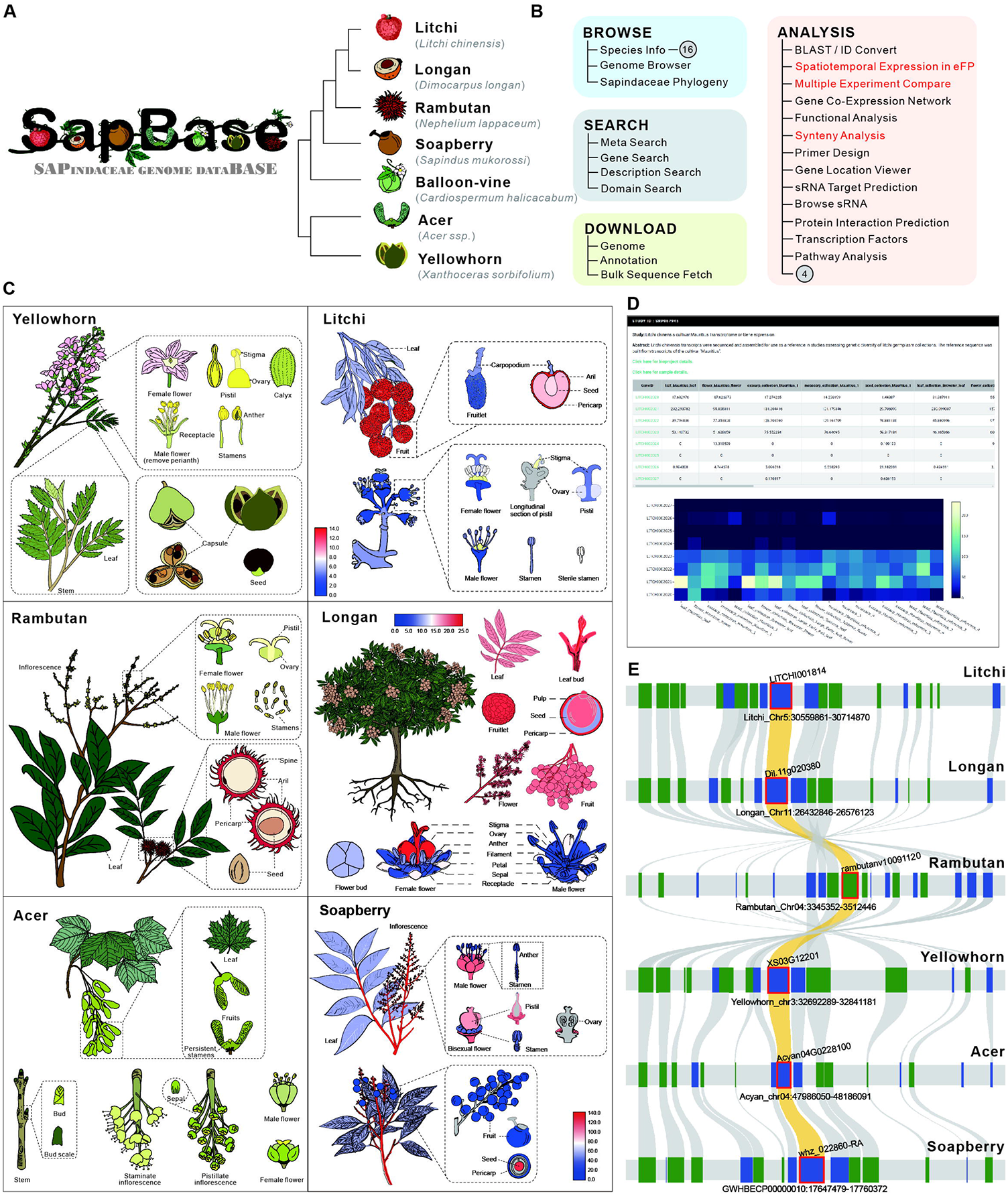
Features of Sapinaceae Genomic DataBase (SapBase). (A) List of current species with genomic data hosted in SapBase. (B) Structure of main functions in SapBase. Numbers in circles show denotes the species number or the number of other functions that cannot be shown here due to space limit. Functions highlighted in red are the three selected functions used for demonstration. (C) eFP (electronic Fluorescent Pictograph) used for gene expression presentation of six Sapinaceae species: Yellowhorn, Rambutan, Acer, Litchi, Longan and Soapberry. The former three are in their original form without displaying expression data while the latter three are in their heatmap form to show the expression of certain genes. Red color denotes a high expression level of gene expression while blue color corresponds to a low expression level. All eFP graphs in SapBase are interactive to view spatial expression of genes. (D) A representative screenshot from the Multiple Experiment Compare function. Expression levels of several genes are listed in a table and displayed in an interactive heatmap. (E) An exemplative result generated from the Synteny Analysis function, using the gene (“LITCHI001814”) as the input.

## Search Strategy

As a multifunctional resource hub, SapBase provides a set of search strategies. Starting from a simple gene identifier, users can obtain its functional annotation, gene structure annotation, domain annotation, and sequences. BLAST, the most commonly used sequence search engine, has been integrated into SapBase for quick nucleic acid or protein sequence comparison among species of interest. A practical ID Convert function is provided for mapping genes from Sapindaceae species to their best homologs in most-researched plant species, such as Arabidopsis, rice, tomato, etc. To maximize the search capacity of SapBase, we designed a sophisticated “Meta Search” module, which provides a “Google-like” search function. Users are allowed to search using any related information, such as gene identifiers, gene function descriptions and even DNA/protein sequences, and SapBase will automatically identify the input content, carry out data search, and return the best matching results.

## Data Analysis

Aside from a datahub, we also aim to develop SapBase as a powerful analytic platform. Currently, a great variety of analysis functions are available in SapBase (Fig.1B). Spatiotemporal Expression shown in eFP (electronic Fluorescent Pictograph) is a feature function designed to intuitively visualize gene expression in customized pictographic representations of plants. We have constructed eFP profiles for all the seven plants (Fig. 1C). There are two other entries in the Gene Expression module: Multiple Experiment Comparison and Co-expression. The former accommodates transcriptional expression profiles generated from all publicly available RNAseq datasets of Sapindaceae plants (Fig. 1D), which can be used for quick assessment of the expression changes of ideal genes under different experimental conditions. The Co-expression function, based on the expression profiles, is implemented to integrate all co-expression gene networks, where users can easily obtain the co-expressed and interconnected counterparts by simply entering the identifier of a gene of interest.

SapBase also provides functions for comparative genomic analyses. Synteny Analysis can be used to quickly explore the evolution and diversification of large syntenic gene blocks (Fig. 1E), and Homology Find function allows users to quickly obtain the optimal homologous gene set for the genes of interest. In addition, a pack of other functions is incorporated in the SapBase, from various integrative data analysis pipelines like Gene Function Enrichment, Gene Pathway Analysis, and Protein Interaction Network, to out-of-the-box tools, including Gene Location Viewer, small RNA Target Prediction, PCR Primer Design, etc.

## Others

SapBase provides entries for downloading all Sapindaceae genomic data and resources, including raw sequencing data, genome sequences, gene annotations, and expression matrices. Users can use the “Bulk Data Fetch” to grab sequences for a large number of genes or chromosome regions in a batch mode. A well-curated list of state-of-the-art software, related genomic databases, or web servers are also recommended on the RESOURCE page.

## Conclusion

By collating publicly released genomes and omics data for seven Sapindaceae species, we have developed SapBase, which provides a one-stop-shop for all Sapindaceae genomic resources, ensuring a convenient and efficient access and usage of all these resources for daily research. As a long-term development project, SapBase will be continuously maintained and updated as a central datahub and analytic platform for researchers working on Sapindaceae or related areas.

## Acknowledgment

We thank all members of XIALAB and the National Litchi and Longan Industrial Technology Consortium of China for their suggestions and testing of SapBase. We are also grateful for the effort other researchers have devoted to the genomic research of Sapindaceae plants.

## References

Chen, C., Li, J., Feng, J., Liu, B., Feng, L., Yu, X., Li, G., Zhai, J., Meyers, B. C., and Xia, R. (2021). sRNAanno—a database repository of uniformly annotated small RNAs in plants. Hortic. Res. 8:45.

Edger, P. P. (2022). The power of chromosome-scale, haplotype-resolved genomes. Mol. Plant 15:393–395.

Hu, G., Feng, J., Xiang, X., Wang, J., Salojärvi, J., Liu, C., Wu, Z., Zhang, J., Liang, X., Jiang, Z., et al. (2022). Two divergent haplotypes from a highly heterozygous lychee genome suggest independent domestication events for early and late-maturing cultivars. Nat. Genet. 54:73–83.

Liang, Q., Li, H., Li, S., Yuan, F., Sun, J., Duan, Q., Li, Q., Zhang, R., Sang, Y. L., Wang, N., et al. (2019). The genome assembly and annotation of yellowhorn (Xanthoceras sorbifolium Bunge). GigaScience 8:6.

Lin, Y., Min, J., Lai, R., Wu, Z., Chen, Y., Yu, L., Cheng, C., Jin, Y., Tian, Q., Liu, Q., et al. (2017). Genome-wide sequencing of longan (Dimocarpus longan Lour.) provides insights into molecular basis of its polyphenol-rich characteristics. GigaScience 6:5.

Lyu, J. (2022). Two domestication routes intersect. Nat. Plants 8:96.

Xue, T., Chen, D., Zhang, T., Chen, Y., Fan, H., Huang, Y., Zhong, Q., and Li, B. (2022). Chromosome-scale assembly and population diversity analyses provide insights into the evolution of Sapindus mukorossi. Hortic. Res. 9.

Yang, J., Wariss, H. M., Tao, L., Zhang, R., Yun, Q., Hollingsworth, P., Dao, Z., Luo, G., Guo, H., Ma, Y., et al. (2019). De novo genome assembly of the endangered Acer yangbiense, a plant species with extremely small populations endemic to Yunnan Province, China. GigaScience 8:7.

Zhang, W., Lin, J., Li, J., Zheng, S., Zhang, X., Chen, S., Ma, X., Dong, F., Jia, H., Xu, X., et al. (2021). Rambutan genome revealed gene networks for spine formation and aril development. Plant J. 108:1037–1052.

